# Dissociating experience-dependent and maturational changes in fine motor function during adolescence

**DOI:** 10.1101/2021.11.19.469287

**Authors:** Andrea Berencsi, Ferenc Gombos, Patrícia Gerván, Zsófia Tróznai, Katinka Utczás, Gyöngyi Oláh, Ilona Kovács

## Abstract

Adolescence is a sensitive period in motor development but little is known about how long-term learning dependent processes shape hand function in tasks of different complexity. We mapped two fundamental aspects of hand function: simple repetitive and complex sequential finger movements, as a function of the length of musical instrumental training. We controlled maturational factors such as chronological and biological age of adolescent female participants (11 to 15 years of age, n=114). We demonstrated that experience improves performance as a function of task complexity, the more complex task being more susceptible for experience driven performance changes. Overall, these results suggest that fine motor skills involving cognitive control and relying on long-range functional brain networks are substantially shaped by experience. On the other hand, performance in a simple repetitive task that explains fine motor speed is primarily shaped by white matter development driven by maturational factors.

## INTRODUCTION

Appropriate hand function is fundamental for human performance during daily living activities and it is a basic requirement for many occupations beyond that. Fine motor skills are determined both by genetically guided maturational processes and by experience-dependent changes during development. Adolescence is a sensitive period in development when the motor system reaches adult-like features by brain reorganization and musculoskeletal changes (Farr and Khosla 2015). Since hand function is crucial in participation in education and in professional life, it is critical to understand how maturational and long-term learning dependent processes may differentially shape various aspects of fine motor performance. Musical instrumental training has been previously used as the model of long-term experience-dependent changes (Schlaug et al. 1995). Our aim was to map two fundamental aspects of hand function, the simple repetitive and complex sequential finger movements as a function of musical instrumental training while controlling maturational factors in adolescence.

The development of fine motor function in terms of simple and complex finger movements (Dorfberger, Adi-Japha, and Karni 2009; Gervan, Berencsi, and Kovacs 2011) and the maturation of the underlying brain areas continue well into adolescence (Wierenga et al. 2014). It involves synaptic pruning and synaptogenesis in the cortical and subcortical grey matter structures including cerebral cortex, basal ganglia and cerebellum (Foulkes and Blakemore 2018; Vijayakumar et al. 2021). Myelination and microstructural development in the white matter of the brain including the motor related areas is associated with sex steroid levels and pubertal stage and suggest the presence of pre-programmed hormonal maturation driven processes (Herting et al. 2012, 2017). Adolescence is also a period of the transition of functional networks from local recruitment to more distributed networks involving long-range connections between cortical areas and cortical and subcortical structures (Fair et al. 2009; Oldham and Fornito 2019). These extended large-scale connections involve motor related networks (Ciechanski, Zewdie, and Kirton 2017) and also those that account for the cognitive and attentional aspects of a complex motor task such as the cingulo-opercular or control networks (Grayson and Fair 2017; Power et al. 2010).

The maturational processes are paralleled by motor experience from childhood to adulthood, leading to their interaction during development (Dow-Edwards et al. 2019). The learning of finger sequences results in the reorganization of functional neural networks in motor cortical areas (Kami et al. 1995; Karni et al. 1998)). These rearrangements in long-range connections correlate with motor performance improvement in speed and accuracy of fine motor sequences (Karni et al. 1998). Improvement during the learning of motor sequences that involve cognitive control is associated with the close interaction of local motor cortical activity and remote functional activity in the large scale frontoparietal network (Maruyama et al. 2021). Furthermore, professional adult musicians have been studied previously to investigate long-term learning, and musical training has been shown to be accompanied by structural and functional changes in multiple cerebral motor and sensory cortices and the cerebellum (Elbert et al. 1995; Gaser and Schlaug 2003; Lee, Chen, and Schlaug 2003; Schlaug et al. 1995). Experience-dependent changes may also involve motor related white matter structures in the long-term (Lee et al. 2003). On the other hand, the association between the experience-dependent alterations in the cerebral white matter and the amount of changes in motor performance is still contradictory (Lakhani et al. 2016).

Fine motor performance of the hand in terms of independent finger movements is frequently approached by the finger tapping (FT) task. The two subtypes of the FT task show differential correlation with maturational and experience-dependent changes. Performance in the sequential finger tapping task correlates with development (Dorfberger et al. 2009) and also with long-term learning related changes in motor cortical areas (Karni et al. 1998). In a recent study, performance in a finger sequence task was related to brain functional connectivity but not to corticospinal excitability (Herszage et al. 2020). In this type of the task that involves more cognitive control (Wong and Krakauer 2019), improvement during learning was related to the functional connectivity of the frontoparietal network including primary motor cortex (Maruyama et al. 2021). On the other hand, performance in the non-sequential repetitive FT task shows association with development (Dorfberger et al. 2009) and with white matter maturation from childhood to adulthood (Bartzokis et al. 2010). Our goal was to study how long-term fine motor experience shapes these two types of motor performance until the period of adolescence.

Musical instrumental training usually starts at an early age, involves regular daily practice sessions for years, represents a highly specific practice in motor terms and may clearly determine some aspects of fine motor performance. Therefore, the length of musical instrumental training can be applied as a model of long-term fine motor experience. We carefully controlled the factors that may cause bias when examining fine motor function in an adolescent group with a maturational spurt. Therefore, to fully control the possible bias caused by differences in gender (Dorfberger et al. 2009; Skogan et al. 2018) and socioeconomic status (Piek et al. 2008), these factors were kept invariable in our sample. Furthermore, the period of adolescence may be characterized by variability in the level of biological maturation even within a given age group. Therefore, in our study we aimed to control maturation in not only the means of chronological age but also that of biological age. Skeletal age is suggested to be a predictor of biological maturation (Holienka et al. 2017) that in turn can explain the variance in gross motor performance up to 6.1% in boys and 20.4% in girls (Freitas et al. 2018). To our present knowledge, no such relationship has been established in relation to fine motor control. Therefore, skeletal age of participants was additionally measured to control for possible differences in biological maturation.

Our hypothesis was that performance in the more complex sequential fine motor task is susceptible to musical instrumental experience and also shows the previously found maturational changes as a function of age (both chronological and bone age). We further assumed a differential effect on the maximum fine motor speed reflected by the simple repetitive finger tapping task. Our hypothesis was that maximum fine motor speed that is related to white matter development will be associated with maturation in terms of age but less associated with learning dependent changes in terms of the length of musical instrumental experience.

## METHODS

### Participants

The 114 female participants of the study were part of a larger developmental study cohort of the MTA - PPCU Adolescent Development Research Group (Kovács et al. 2021). Participants’ chronological ages ranged from 11.1 to 15.1 years (*M* = 13.1, SD = 1.1), bone age ranged from 11.2 to 16.0 years (*M* = 13.2, SD = 1.2) and their musical instrumental experience ranged from 0 to 8 years (*M* = 3.1, SD = 2.8). In order to control for the impact of education and socioeconomic status, all participants were recruited from top rated secondary schools in Budapest (based on official university enrolment data). Parents’ educational level was also obtained by a screening questionnaire, and 90.4 percent of the parents have a university or college degree. Students were recruited via contacting schools, and advertisements were also placed on Facebook and online magazines. All subjects spoke Hungarian as their first language. Based on the screening questionnaire none of the included subjects has a history of developmental disorder, learning disability or neurological disorder. 16 of them were left-handed based on the reports of hand preferred to perform unimanual tasks.

We excluded one participant who did not report on the length of musical training, one participant who was diagnosed with ADHD, and two participants where measurement equipment failed.

Biological maturity (bone age) was assessed in their school or in the Research Centre for Sport Physiology at the University of Physical Education. The assessment of fine motor movement took place at the Developmental Neuroscience Laboratory of the Institute of Psychology at PPCU. The Hungarian United Ethical Review Committee for Research in Psychology (EPKEB) approved the study (reference number 2017/84). Written informed consent was obtained from all the subjects and their parents. Participants were given gift cards for their attendance.

### Musical instrumental experience

Musical instrumental experience of participants was obtained through a questionnaire. The number of years spent with musical instrumental training was calculated as the summary of playing an instrument for a full year. Number of years of playing an instrument or multiple instruments were maximized in 8 years. There was no restriction regarding the type of musical instrument.

### Bone age measurement

Biological age refers to the biological development of an individual providing also an estimate of the maturity status (advanced, average, delayed). One of the most commonly used methods to estimate maturity is the assessment of bone age, which is a method evaluating the progress of ossification. Body height was measured with a standard anthropometer (DKSH Switzerland Ltd, Zurich, Switzerland) and body mass with a digital scale (Seca). Age at menarche was collected retrospectively by interviewing the participants. Bone age was estimated with an ultrasound-based device (Sunlight Medical Ltd, Tel Aviv, Israel). During the measurement, the ultrasound passes through the subjects left wrist evaluating the distal epiphyseal and diaphyseal ossification of the two forearm bones (radius and ulna). Measurements were repeated five times and the transducers were raised by 2 mm for each measurement. The device estimated bone age (in years and months) by measuring the speed of sound (SOS) and the distance between the transducers. The same person performed all measurements.

### Fine motor tasks

Motor tasks were performed both with the dominant and non-dominant hands.

#### Sequential finger tapping (FT) task

Sequential FT task was a four-element finger-to-thumb opposition task in the order of index-ring-middle-little fingers. Ten sets of 16 repetitions were performed. Participants were asked to perform as fast and as correctly as possible. Data acquisition started when participants were able to repeat three consecutive sequences correctly, eyes closed. Dependent variable was Performance Rate (PR) calculated as the number of finger taps in correctly performed sequence/s. PR mean of ten blocks was used in each participant.

#### Simple repetitive finger tapping task (Index FT task)

The non-sequential task was an index finger to thumb opposition task. 64 taps were performed with maximum speed. Dependent variable was the number of taps/s. Mean of three blocks of 64 taps were calculated in each participant.

#### Data acquisition and processing

Order and timing of finger tapping was detected by a custom-made data glove. It contained metal electrodes on the palmar surface of the distal phalanges on each finger. Data glove was attached to serial-to-USB hub connected to a laptop computer. A custom-made Java based software (Gervan et al. 2011) acquired the data of timing and order of finger taps and also automatically calculated PR and number of index finger taps/s of each participant.

#### Statistical analysis

Pearson’s correlation coefficients were calculated to evaluate the association between musical experience, age and bone age on sequential and repetitive finger tapping performance. Partial correlations controlled for age and bone age were calculated to reveal the effect of experience, chronological age and bone age, respectively. Significance level was set at p<.05.

## RESULTS

### Sequential finger tapping task

Results of the Pearson correlation indicated that there was a significant positive association between musical instrumental experience and PR on the non-dominant hand (r(114) = .538, p = 6.4889E-10, 95% CI [.384, .667]) and PR on the dominant hand PR (r(114) = .451, p = 4.6769E-7, 95% CI [.278, .602]; Fig. 1.a-b.). Chronological age was also positively correlated with PR on the non-dominant hand (r(114) = .361, p = .000078, 95% CI [.177, .522]) and dominant hand (r(114) = .391, p = .000017, 95% CI [.203, .576]). Bone age showed positive association with PR on non-dominant (r(114) = .291, p = .001712, 95% CI [.128, .439]) and dominant hand (r(114) = .379, p = .000033, 95% CI [.204, .538]). Chronological age and bone age were also significantly positively correlated (r(114) = .758, p = 1,5834E-22, 95% CI [.695, .812]).

**Figure 1.**
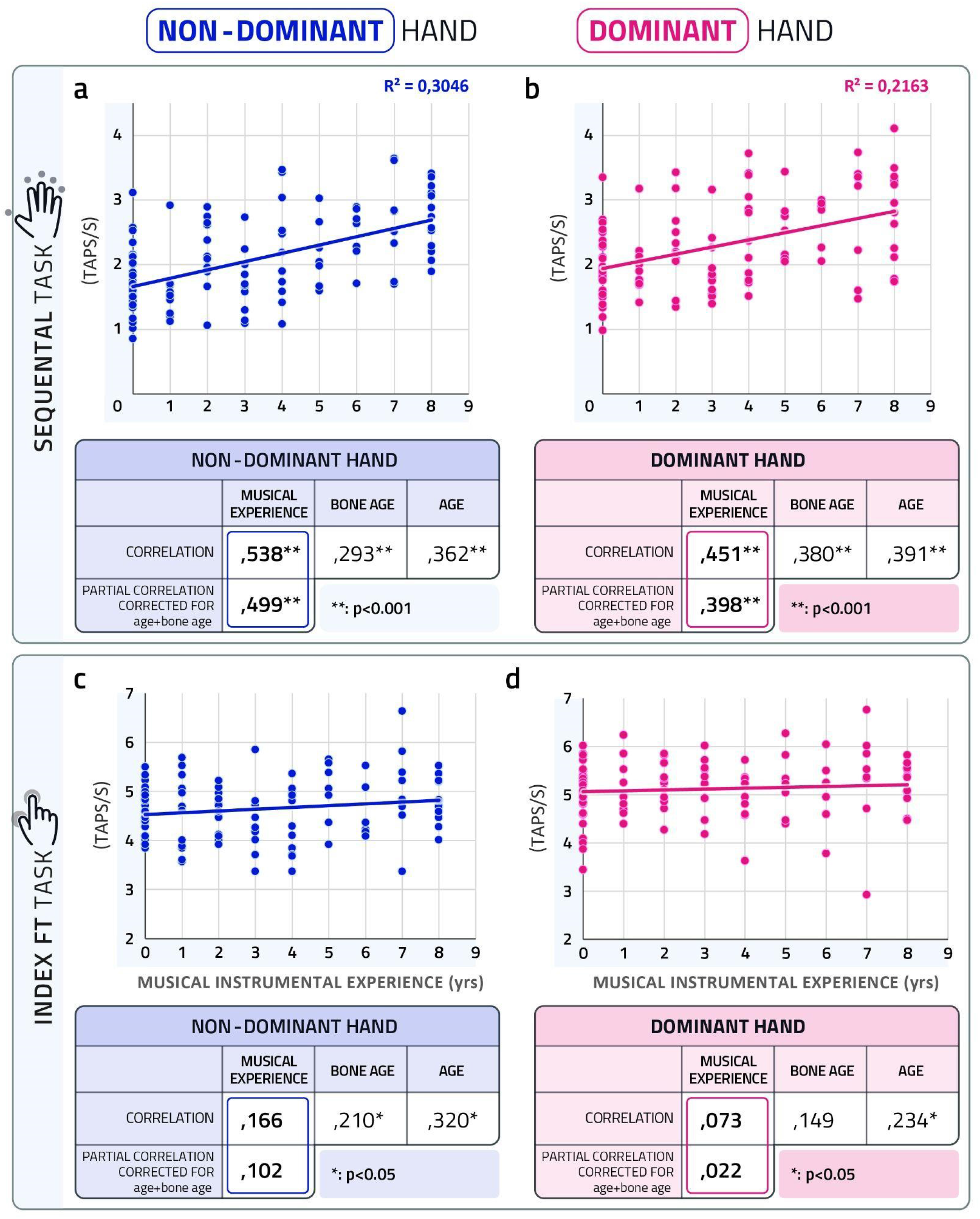
Dissociation of experience-dependent changes in fine motor performance as a function of years spent in musical training in sequential and repetitive finger movements. Length of instrumental training significantly correlates with sequential performance improvement both in non-dominant (a) and dominant (b) hand. Simple repetitive finger tapping task performance is not related to musical experience either in non-dominant (c) or dominant (d) hand. After correction for age and bone age, sequential motor performance still strongly correlates with musical training (a and b), possibly reflecting the long-term musical instrumental experience dependent changes in motor cortical areas and related long-range networks. After correction, index finger tapping task exhibit no significant partial correlation with instrumental training (c and d). It likely reflects that experience dependent changes due to musical instrumental training have no significant effect on myelination related function of index finger tapping.

When Pearson correlation was controlled for chronological age and bone age, significant positive association between musical instrumental experience and PR remained high both in the dominant (r(110) = .398, p = .000014, 95% CI [.234, .546]) and the non-dominant hand (r(110) = .500, p = 1.9685E-8, 95% CI [.352, .627]). PR was positively correlated with chronological and bone age when correlation was controlled for musical experience both in the dominant (r(111) = .329, p = .000368, 95% CI [.170, .486] and r(111) = .322, p = .001, 95% CI [.170, .486]) and non-dominant (r(111) = .287, p = .002, 95% CI [.135, .453] and r(111) = .211, p = .025, 95% CI [.037, .380]) hands, respectively. Correlation controlled for musical instrumental experience and bone age showed significant positive association between chronological age and PR in the non-dominant (r(110) = .200, p = .034, 95% CI [.036, .336]) but not on the dominant hand (r(110) = .142, p = .136, 95% CI [-.039, .319]). Correlation controlled for musical instrumental experience and chronological age revealed no association between bone age and PR neither in the dominant (r(110) = .121, p = .202, 95% CI [-.061, .301]) or the non-dominant hand (r(110) = -.005, p = .954, 95% CI [-.169, .166]). In summary, the performance in a sequential fine motor task is associated with developmental factors in terms of both chronological age and bone age, but the effect of bone age independent from chronological age is not present. On the other hand, the independent effect of long-term motor experience due musical training explains the majority of the variance in the complex fine motor task.

### Repetitive finger tapping task

Results of the Pearson correlation indicated no significant association between musical instrumental experience and index finger tapping speed on either the non-dominant hand (r(114) = .166, p = 0,078, 95% CI [-.019, .333]) or the dominant hand (r(114) = .073, p = .439, 95% CI [-.145, .282]; Fig. 1. c-b.). On the other hand, chronological age showed positive association with index FT speed on the non-dominant hand (r(114) = .320, p = .000494, 95% CI [.142, .491]) and the dominant hand (r(114) = .234, p = .012, 95% CI [.045, .411]). Bone age also positively associated with the index finger FT speed on the non-dominant (r(114) = .210, p = .025, 95% CI [.050, .376]) hand but not with that of the dominant hand (r(114) = .149, p = .115, 95% CI [-.022, .324]).

After controlling for age and bone age, association between index finger tapping speed and musical instrumental experience has diminished both in the dominant (r(110) = .022, p = .815, 95% CI [-.174, .215]) and the non-dominant hand (r(110) = .102, p = .284, 95% CI [-.083, .272]). These results suggest that age and maturation have all-decisive roles on simple repetitive fine motor performance in adolescence. When correlation was controlled for musical experience, index finger tapping was positively correlated with chronological age in the dominant (r(111) = .225, p = .017, 95% CI [.029, .401]) and non-dominant hand (r(111) = .295, p = .002, 95% CI [.108, .475]) but not with bone age in the dominant (r(111) = .136, p = .150, 95% CI [-.040, .291]) and non-dominant hand (r(111) = .180, p = .056, 95% CI [.003, .351]). Correlation controlled for musical instrumental experience and bone age showed significant positive association between chronological age and repetitive finger tapping performance in the non-dominant (r(110) = .244, p = .009, 95% CI [.071, .416]) and the dominant hands (r(110) = .187, p = .048, 95% CI [.006, .368]). Correlation controlled for musical instrumental experience and chronological age revealed no association between bone age and index finger tapping performance neither in the dominant (r(110) = -.048, p = .612, 95% CI [-.207, .134]) or the non-dominant hand (r(110) = -.061, p = .520, 95% CI [-.210, .101]).

## DISCUSSION

We investigated two aspects of fine motor performance by employing a sequential and a repetitive finger tapping task to reveal the effect of long-term motor experience in terms of the length of musical instrumental training. The results showed a differential effect of musical instrumental training in the repetitive and sequential tasks. While performance in the more complex sequential task seems to be strongly affected by the amount of experience in musical instrumental training, the same effect is lacking in the simple repetitive task.

Experience-dependent effect on sequential task in terms of the length of musical instrumental training is in line with our hypothesis. We assumed that performance in the more complex sequential fine motor task is susceptible to musical instrumental experience. A sequential component in a motor task requires cognitive control and the coordinated interaction of several distant brain areas as compared to the simple repetitive task such as the dorsal premotor cortex, superior and posterior parietal areas and posterior cingulate cortex (Haaland et al. 2004; Hamano et al. 2020; Yokoi and Diedrichsen 2019). This cooperation depends upon the long-range networks between distinct cortical areas and between cortical and subcortical areas (Asato et al. 2010) that mature late during adolescence, especially between the late maturing functional hubs of the frontal and the parietal cortices (Oldham and Fornito 2019).

The phenomenon of a lacking impact of musical experience on a simple repetitive task in non-professional adolescent participants in our study suggests that maximum fine motor speed is not determined by musical experience. Moreover, regardless of the amount of musical instrumental practice our participants’ results also correspond to the recently updated norms of finger tapping performance (Skogan et al. 2018). Similar results were reported in a study of adult professional piano players where neither the age of musical training initiation nor the amount of musical training before the age of twenty was a predictor of repetitive piano keystrokes (Furuya et al. 2015). The development of maximum motor speed as a function of repetitive finger tapping shows an inverted U-shaped curve peaking at 38 years and is related to the white matter myelination level in males (Bartzokis et al. 2010). The microstructure of callosal connections and corticospinal tracts is suggested to develop early and have a relative stability across ages from adolescence to adulthood (Slater et al. 2019). It may be reflected in the transition from a rapid to a slower development in index FT performance that reached between puberty and young adulthood (Fietzek et al. 2000; Gasser et al. 2010) with a continuous improvement in index FT speed in adolescence (Dorfberger et al. 2009). In spite of the ongoing myelination and age-related changes in the simple repetitive task during adolescence, long-term highly specific fine motor training did not interact with this process on the performance level.

Our results showed the effect of maturation in terms of correlation with chronological age on both types of finger tapping performance. This effect is long known in the field (Dorfberger et al. 2009; Gervan et al. 2011; Skogan et al. 2018) and was therefore also assumed to appear in the present study. On the other hand, the independent effect of biological maturation from chronological age in terms of bone age on fine motor performance was not found in our study. It should be noted that there was a high positive correlation between chronological age and bone age and this association should be there in any such study. Therefore, the lack of independent impact in terms of biological age may in part be due to these phenomena in our study. To our knowledge, there is no previous study that took biological and chronological age into account simultaneously when controlling for developmental effects in fine motor performance. A study to explore the relationship between fine motor performance and bone age did not reveal such effect in 5-9 years old boys and girls (Kerr 1975) while chronological age was not controlled in this regard.

Our results clearly demonstrated that for complex movements involving higher cognitive demands such as advanced planning, composition of sequences and the cooperation of long-distance brain areas, the effect of experience overrides the effect of age and maturity until adolescence in typically developing populations. It suggests that late maturing long-range brain networks may be more susceptible to prolonged experience dependent changes. On the other hand, performance in simple repetitive movements is driven by maturation, and age rather than experience determines the maximum motor speed of the hand. These results have several practical implications. From the educational point of view, while maximum motor speed may be a personal characteristic associated with age, production of skilful complex movements may be a function of the amount of practice spent with fine motor training in the typically developing population. While other types of motor skills certainly need to be also mapped in this regard, its applications may be useful in educational settings and during career planning before adolescence. Furthermore, the present phenomena should be further examined in atypical development since there are neurodevelopmental disorders such as autism spectrum disorder (Oldehinkel et al. 2019), attention deficit hyperactivity disorder (Bos et al. 2017) and rare disorders (Gombos, Bódizs, and Kovács 2017) that involve both atypically developing long-range brain networks including motor subsystems and altered hand function (Berencsi, Gombos, and Kovács 2016; Rommelse et al. 2007; Travers et al. 2017).

Our study’s strength causes its limitation at the same time. While we carefully controlled the inclusion criteria regarding gender, age, education and socioeconomic status it may reduce the generalizability of the results. On the other hand it was necessary to exclude as many confounding factors as possible when looking at long-term effects in a cross sectional design.

In summary, our results demonstrated the dissociation of the white matter maturation related repetitive motor task and the long-range network associated complex sequential fine motor tasks’ susceptibility to prolonged practice. Complex fine motor performance is shaped by experience dependent changes in contrast to maturation determined repetitive task.

## Notes

### Competing Interest Statement

The authors have declared no competing interest.

